# Single-cell transcriptomics defines an improved, validated monoculture protocol for differentiation of human iPSCs to microglia

**DOI:** 10.1101/2022.08.02.502447

**Authors:** Sam J Washer, Marta Perez-Alcantara, Yixi Chen, Juliette Steer, William S James, Andrew R Bassett, Sally A Cowley

**Affiliations:** James and Lillian Martin Centre for Stem Cell Research, Sir William Dunn School of Pathology, University of Oxford, South Parks Road, Oxford, OX1 3RE; Wellcome Sanger Institute, Wellcome Genome Campus, Hinxton, CB10 1SA; Open Targets, Wellcome Genome Campus, Hinxton, CB10 1SA

**Keywords:** Microglia, differentiation, protocol, induced pluripotent stem cells, Single cell RNA sequencing

## Abstract

There is increasing genetic evidence for the role of microglia in neurodegenerative diseases, including Alzheimer’s, Parkinson’s, and motor neuron disease. Therefore, there is a need to generate authentic *in vitro* models to study human microglial physiology. Various methods have been developed using human induced Pluripotent Stem Cells (iPSC) to generate microglia, however, systematic approaches to identify which media components are actually essential for functional microglia are mostly lacking. Here, we systematically assess medium components, coatings, and growth factors required for iPSC differentiation to microglia. Using single-cell RNA sequencing, qPCR, and functional assays, with validation across two labs, we have identified several medium components from previous protocols that are redundant and do not contribute to microglial identity. We provide an optimised, defined medium which produces both transcriptionally and functionally relevant microglia for modelling microglial physiology in neuroinflammation and for drug discovery.

## Introduction

Microglia are the resident macrophages of the brain parenchyma, responsible for a broad range of homeostatic functions, including clearance of apoptotic cells and misfolded proteins, and synaptic pruning during neurodevelopment (1). Microglia are associated with a number of neurodegenerative disorders, notably Alzheimer’s Disease (AD) (2–4), Parkinson’s disease (PD) (5), and motor neuron disease/amyotrophic lateral sclerosis (ALS) (6), with many of the genes with highest genetic risk score being expressed in microglia, including; *TREM2, CD33, APOE, LRRK2,* and *C9orf72*.

Studies of primary microglia have been mostly restricted to rodent models, as access to fresh primary human microglia is severely limited. Furthermore, microglia are highly sensitive to their environment, so the removal and culturing of microglia away from their original homeostatic environment is likely to result in immune activation, and they will therefore differ from their *in vivo* state, such that they may not recapitulate the disease cellular physiology *in vitro* (7).

As an alternative to rodent and primary microglia, several labs have developed methods for generating human microglia from induced Pluripotent Stem Cells (iPSC) (8–17) (**TABLE 1**). These protocols all aim to roughly recapitulate the sequence of events in the developing embryo. This approach potentially facilitates the study of the effect of disease genotype on microglial phenotype, as iPSCs provide a limitless resource, faithfully reproducing the donor’s genetic background, which can also be readily gene-edited as required. However, issues with iPSC-based methodologies exist, including: variable reproducibility of protocols between different labs; variation across different cell lines; unchallenged adoption of medium components from other protocols; and use of undefined medium components. The multiple iPSC microglial protocols that have been published (including by our laboratory (10)), use different combinations of growth factors and media supplements including IL-34, TGF-β1, M-CSF, GM-CSF, CD200, CX3CR1, supplements B27 and N-2, and different substrates, in an attempt to recapitulate primary microglia *in vitro*. Several studies require further differentiation steps via co-culture, integration into organoids or xenotransplantation into mice, thus limiting accessibility of these models (12,16). Relevant here, the medium developed by our lab (10) was optimised for co-culture with iPSC-cortical neurons, rather than monoculture. Systematic comparison of the different components has been minimal, to identify which, if any, are redundant or provide minimal benefit to producing iPSC-microglia in monoculture.

**Table 1:**
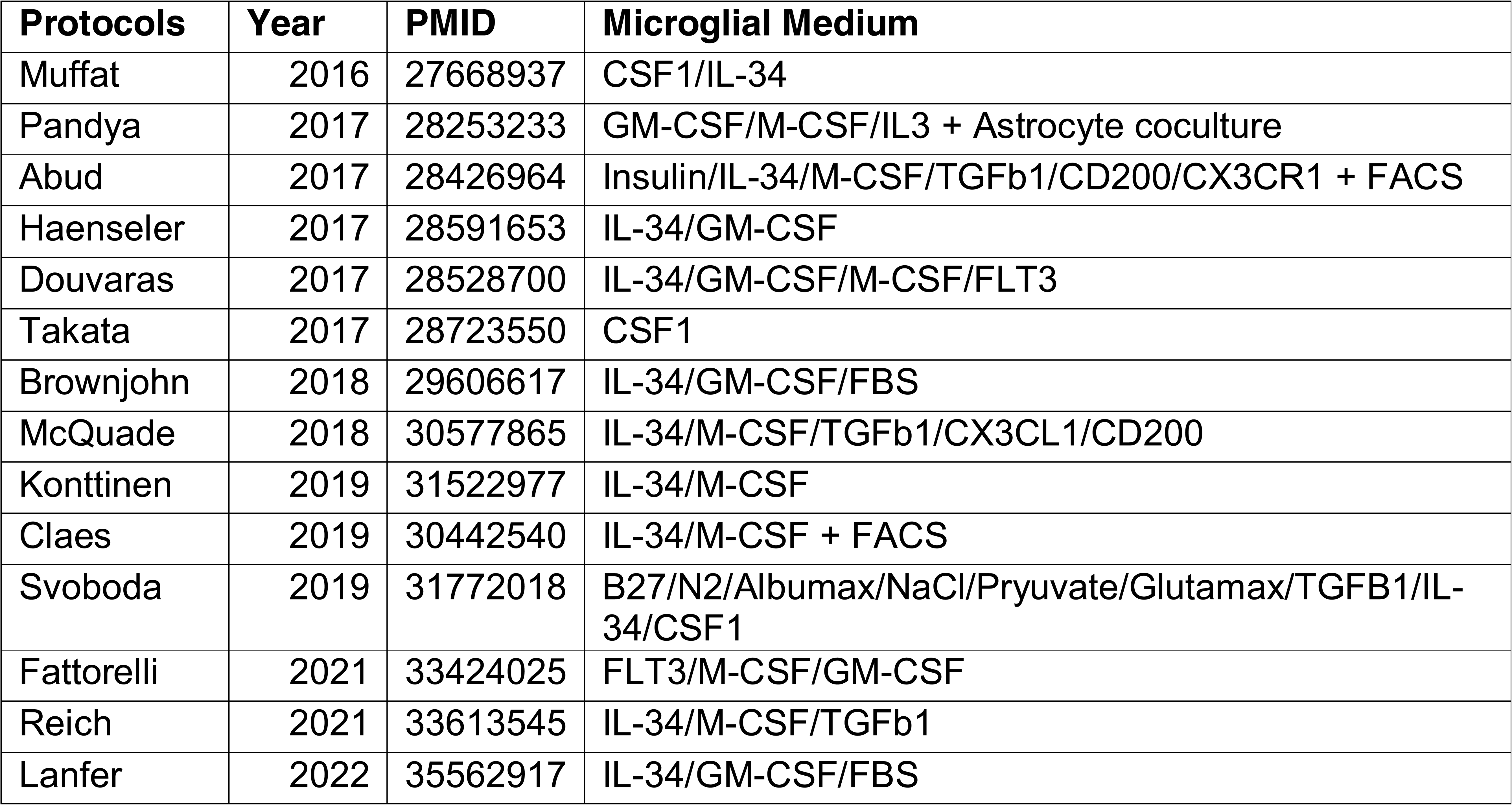
Published medium for differentiating iPSC to microglia.

Here, we set out to systematically compare and identify how different growth factor, medium component and substrate combinations affect resultant iPSC-microglial phenotypes, transcriptomes, and survival in monoculture, in an attempt to identify a medium which can best recapitulate human microglia *in vitro* without the need for co-culture or xenotransplantation. We validated the results across two laboratories, using the same iPSC, basal medium, growth factors, and reagent providers. Building from the previously published microglial differentiation protocol by (10),Haenseler et al. which starts with iPSC differentiation to mesoderm via embryoid bodies, hemogenic endothelium induction, then bulk myeloid differentiation to primitive macrophage/microglia precursors, in a highly scalable pipeline. However, Haenseler et al. optimised the final iPSC-microglia maturation medium for co-culture of iPSC-microglia with iPSC-neurons. Here, we directly compared the addition and removal of key components from this and other published protocols, and assessed their ability to recapitulate human microglia in monoculture *in vitro*. We identify substantial variation between previously published protocols for generating iPSC-microglia and present an optimised media composition which is reproducible across multiple genetic backgrounds, improves microglial retention and transcriptionally most closely corresponds to primary microglia. This iPSC-microglia maturation protocol can be applied to study microglia-associated disease settings and can be scaled for drug and phenotypic screening.

## Methods

### Consent for use of Human Material

All iPSC were obtained with informed consent and with the approval of all the relevant institutions. iPSC lines were generated previously as part of the HipSci project (REC Reference 09/H0304/77, covered under HMDMC 14/013 and REC reference 14/LO/0345), or the EU IMI project, StemBANCC (REC Reference 10/H0505/71) (18).

### iPSC lines

The three lines (all from healthy donors) used in this study form part of the HipSci or StemBANCC initiatives and are listed in (**Table S1**).

KOLF2.1S was generated from the parental KOLF2-C1 line (https://hpscreg.eu/cell-line/WTSIi018-B-1) itself a clonal isolate of KOLF2 (https://hpscreg.eu/cell-line/WTSIi018-B) by correction of a heterozygous loss of function mutation in ARID2 using CRISPR based homology directed repair. Cells were nucleofected (Lonza) with recombinant enhanced specificity Cas9 protein (eSpCas9_1.1), a synthetic sgRNA (target site AAAAGATCACTTGCTAATGC CGG, Synthego) and a single-stranded DNA oligonucleotide homology directed repair template (5’-CACTCTCCTATCAAATGAAAGCAAGCACGTCATGCAACTTGAAAAAGATCCTAAAATCATCACTTTACTACTTGCTAATGCCGGGGTGTTTGACGACAGTAAGTTTTAAGCTGAATGTA-3’, IDT Ultramer) followed by clonal isolation (19). Repaired clones were identified using PCR across the edited region, and sequencing of the amplicons by high throughput sequencing (Illumina MiSeq) and validated by Sanger sequencing. These lines were generated by the Cellular Operations Gene Editing team at the Wellcome Sanger Institute.

SFC841-03-01 (20) and SFC856-03-04 (10) were generated at the James and Lillian Martin Centre for Stem Cell Research, University of Oxford, and are available from EBiSC.

### iPSC culture

All differentiation reagents are listed in **(Table S2)** and all medium compositions are listed in **(Table S3)**. iPSC were cultured in ‘OxE8’ medium (21) (based on the published E8 medium formulation (22)) on Geltrex (Gibco A1413302) coated tissue culture dishes and passaged at 80% confluency with 0.5mM EDTA. Cells were incubated at 37°C, 5% CO_2_ and fed daily with fresh OxE8 medium. HipSci lines were cultured in Essential 8 medium (Gibco A1517001) on vitronectin (Gibco A14700) coated tissue culture wells.

### Differentiation to microglia precursors

iPSC were differentiated into primitive macrophage/microglia precursors as described previously (23). In brief, an Aggrewell 800 plate (STEMCELL Technologies, no. 34815) was prepared by addition of 0.5ml of Anti-Adherence Rinsing Solution (STEMCELL Technologies, no 07010) and centrifuged at 3,000 x g for three minutes to remove bubbles from the microwells. Rinse solution was then removed and the wells washed with 1ml PBS before the addition of 1ml of 2x concentrated EB medium (1x EB medium: OxE8 medium supplemented with 50ng/ml BMP4 (PeproTech no PHC9534), 50ng/ml VEGF (PeproTech no PHC9394), and 20ng/ml SCF (Miltenyi Biotec no 130-096-695)) supplemented with 10µM Y-27632 ROCK inhibitor. iPSC were cultured to 70-80% confluency before washing with 1ml PBS then incubation in TrypLE Express (Gibco, no 12604013) for 3-5 minutes at 37°C, 5% CO_2_. Cells were then lifted and transferred to a falcon tube containing PBS to prepare a single cell solution. Cells were counted and pelleted by centrifugation at 400 x g for five minutes. iPSC were then resuspended in OxE8 with 10µM Y-27632 ROCK inhibitor at a concentration of 4 x 10^6^ cells/ml and 1ml of cells were added to one well of the Aggrewell 800 containing 2x EB media. The Aggrewell plate was then spun at 100 x g for 3 minutes with no braking to encourage even distribution of cells. Aggrewells were incubated for six days at 37°C, 5% CO_2_ with daily feeding of 75% media change with EB media. After six days EB were removed from the plate using a 5ml stripette and transferred to a 40µm cell strainer, remaining EB were flushed from the well with 2x 2ml PBS washes. Dead cells were washed from the EB in the strainer using 4ml of PBS. The filter was then inverted over a six well tissue culture plate and EB washed off using 4ml of differentiation media. EB were then divided evenly across two six well plates before being transferred to two T175 flasks containing 18ml of differentiation media (total medium 20ml). These “Factories” were incubated at 37°C, 5% CO_2_ with weekly feeding of 10ml differentiation media until macrophage/microglia precursors (PreMac) cells started to appear in the supernatant. After this point, usually 5 weeks into differentiation, 50% of media was harvested from the factories and PreMac cells were collected by centrifugation at 400 x g for five minutes. Media removed was replaced with fresh Factory media. Cells were used immediately for experiments or stored for use later in a 1L bioreactor (Corning) containing Factory media spinning at a constant speed of 30rpm as described by (24).

For 19 HipSci lines, iPSCs were dissociated with Accutase (LifeTech A1110501) for 5 minutes at 37°C, 5% CO_2_. Cells were pelleted and resuspended in Essential 8 (Gibco) with 10µM Y-27632 (Stemcell Technologies 72304) and strained through 40µM Pluristrainers (Cambridge Bioscience 43-10040-40) to obtain single cell suspensions. HipSci lines were counted and pooled in equal proportions in Essential 8 with 10µM Y-27632 at a concentration of 200,000 cells/mL. The pooled cell suspension was added in equal volume to 2x EB media for a final concentration of 100,000 cells/mL and transferred to a sterile reservoir. 100µL cell suspension containing 10,000 cells was added to each well in Corning CoStar Ultra Low Cluster round bottom ULA 96-well plates (Corning 7007). The ULA plates were centrifuged at 300 x g for 3 minutes and incubated at 37°C, 5% CO_2_ until Day 3. The 96-well plates were fed with 100µL per well EB media after removing 50µl per well spent media at day 3. The plates were incubated at 37°C, 5% CO_2_ until day 6 when the EBs were transferred using a multichannel pipette with wide orifice tips into a sterile reservoir. The EBs were washed as described above and transferred to gelatin (Sigma G1890) coated T75 flasks (Corning 430641U) in a total of 15mL Factory media. These “Factories” were maintained at 37°C, 5% CO_2_ with total volume incrementally increased over the first 3 feeds from 15mL to the maximum volume of 50mL, followed by bi-weekly 60-80% media change. Around 3 weeks after Factory set up, spent media harvested at each feed contained PreMac cells. To start microglial differentiation, the spent media was passed through 40µm cell strainer (Falcon 352340) and PreMac cells were collected by centrifugation at 400 x g for 5 minutes.

### Maturation to microglia

Microglial precursors were plated in 6 or 12-well plates at a density of 100,000 cells/cm^2^. For Geltrex coating, plates were incubated with Geltrex (Thermofisher, A1413202) for 1hr at 37°C, 5% CO_2,_ before removal and direct addition of cells. Where fibronectin (Sigma, F0895) was used, it was diluted to 10µg/ml in PBS and plates were incubated for 3hr at room temperature, removed, and wells washed three times with water before adding cells. Factors used for the experiments and concentrations are outlined in **(Table S4)** with the following acronym when included in a medium: I (IL-34), T (TGF-β1), M (M-CSF), G (GM-CSF), C (CD200/CX3CL1), B (β-mercaptoethanol), N (N-2), F (FBS). 50% media change was performed every three to four days and maturation was allowed to continue for a total of 14 days. For media containing CD200/CX3CL1, these were added at day 10 of the maturation, as described in (12). An overview of the protocol is provided in (**Figure 1a**). The triaging of factors in the test microglial media, including selection criteria, is shown in **Figure 1b**, exclusion criteria are listed in **Figure 1c** and the final comparison is shown in **Figure 1d**.

**Figure 1:**
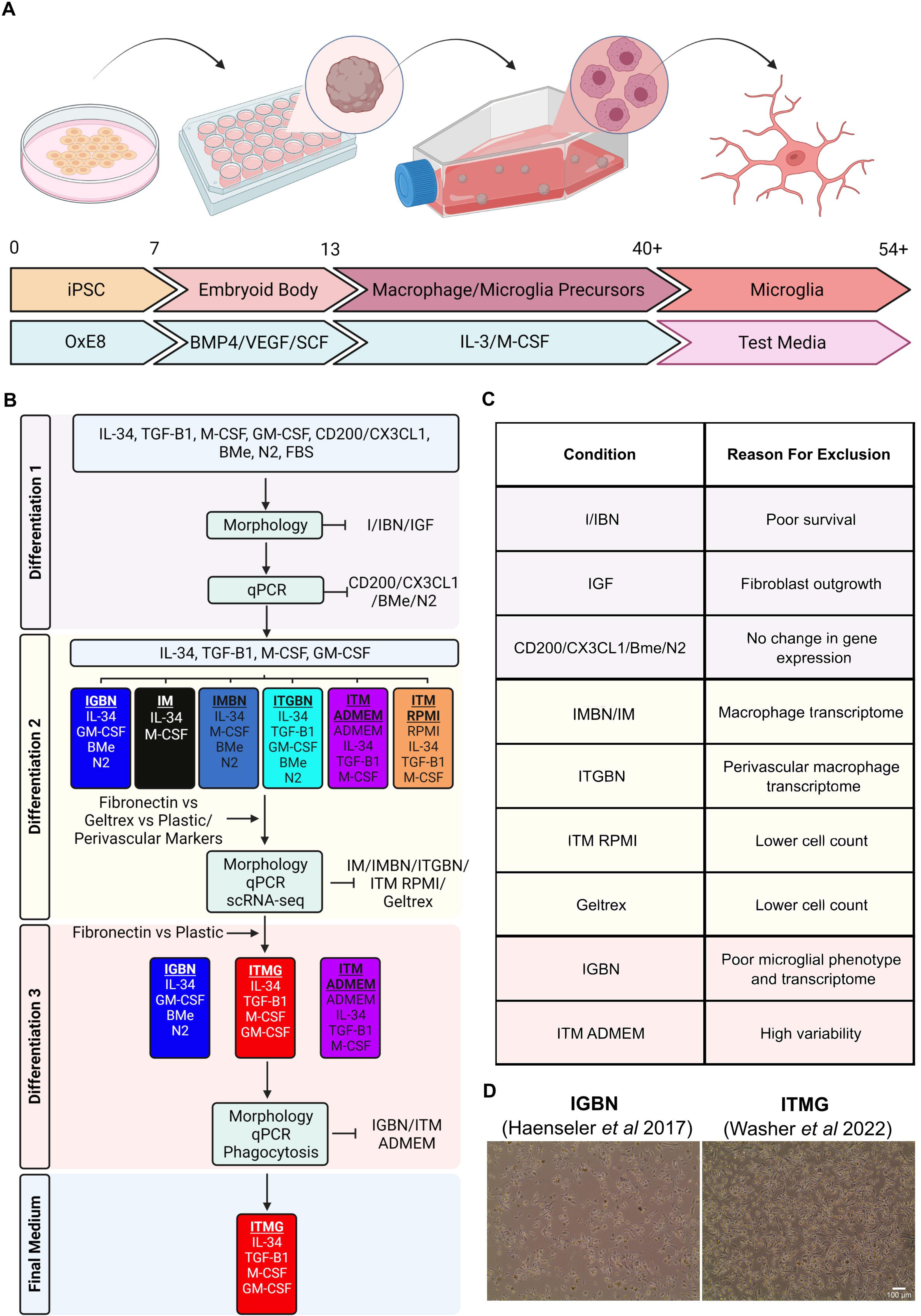
A flowchart of the differentiation process from iPSC to Microglia as described by Haenseler et al. **A)** iPSC are cultured in OxE8 medium for 7d before differentiation into embryoid bodies in Aggrewell plates for 6d in embryoid body medium (BMP4/VEGF/SCF). After 6d embryoid bodies are transferred to T175 flasks and cultured in myeloid differentiation media of IL-3 and M-CSF. At approximately d40 these factories produce microglial precursor cells which are harvested and plated in microglial media and differentiated for 14d. This final step is the target for optimisation in this manuscript. iPSC; induced pluripotent stem cell, EB; embryoid body, SCF; stem cell factor, BMP4; bone morphogenetic protein 4, VEGF; vascular endothelial growth factor, IL-3; Interleukin 3, M-CSF; macrophage colony stimulation factor, d; day. **B)** A flowchart describing the systematic identification of factors required for iPSC-microglial differentiation. After the first triage of 15 media combinations, the morphology analysis removed three conditions due to poor differentiation. Following qPCR analysis four factors could be removed from further medium development. The second set of differentiations focused on IL-34, TGF-β1, M-CSF, and GM-CSF as these were shown to have an effect in the first round of differentiations. Several conditions were removed due to poor microglial identity following scRNA-seq and morphology analysis. In the final differentiation, we identified that our new medium (ITMG) produced iPSC-microglia with a microglial-like transcriptome, improved survival, and performed similarly to our previous medium in functional assays. **C)** Conditions and media tested and their reason for exclusion. **D)** Representative images of d14 microglia cultured in our previous media (IGBN) and the media presented in this manuscript (ITMG). Scale bar 100µm.

### RNA extraction

RNA was extracted from cells using TRIzol LS (Invitrogen, 10296010) and purified using the Zymo DirectZol RNA Miniprep Kit (Zymo Research, R2050) following manufacturer’s instructions. Briefly, media was removed from the microglia and wells washed with PBS, 250µl TRIzol LS reagent was then added to each well and cells flushed off using a p1000 pipette. Lysates were collected into a 1.5ml tube and either frozen at −80°C or immediately extracted following the Zymo DirectZol protocol. RNA was eluted in 25µl RNase/DNase free water and RNA concentrations quantified by Nanodrop.

For HipSci pooled differentiations, RNA was extracted using AllPrep DNA/RNA Mini kit (QIAGEN 80204) or RNeasy Mini kit (QIAGEN 74104) with QIAshredder (QIAGEN 79654) and treated with Turbo DNase (Life Tech AM1907) following manufacturer’s instructions.

### qPCR

Maximal RNA was converted to cDNA using the High Capacity RNA-cDNA conversion kit (Applied Biosystems, 4387406) following manufacturer’s instructions. Assuming a 100% conversion rate, cDNA was then diluted to 1ng/µl using RNase/DNase free water for use in qPCR. qPCR reactions were undertaken in triplicate using Power SYBR Green PCR Master Mix (Applied Biosystems, 4367659) following manufacturer’s instructions in 384 well plate format with the QuantStudio 5 system. For each cDNA conversion a qPCR for housekeeping gene panel was undertaken to confirm the most stably expressed housekeeper to use for the marker analysis. Primers are listed in **(Table S5).** GeNorm was used to analyse housekeeper stability and the two most stable housekeepers were taken forward for use in the marker qPCR panel (25). We undertook qPCR on a panel of known microglial and perivascular marker genes. Data was analysed in the statistical package R using linear regression, multiple testing of p-values were corrected by Bonferroni. Raw Ct values were collected from the software, SDs of the triplicates were calculated, any outliers (determined by ±2.5 SD about the mean) were excluded and the mean Ct value was recalculated. Differential expression was calculated using the 2^−ΔΔCT^ method using the selected housekeepers as normalising controls (26). The baseline condition is defined in each experiment as either RNA extracted from cells in the baseline microglial media (IGBN) or from microglial precursors.

### Single cell RNA sequencing

To harvest cells for single-cell RNAseq, the cells were incubated in a dissociation buffer of DPBS with 5mM EDTA (Invitrogen 15575-038) and 4mg/mL lidocaine hydrochloride monohydrate (Sigma L5647) at 37°C for 15 mins. An equal volume of 0.04% BSA in DPBS was added to each well to avoid sticking of cells to vessels. Cells were collected into centrifuge tubes pre-coated with 0.04% BSA and centrifuged at 400 x g for 5 mins. The pellet was resuspended in 0.04% BSA and strained through MACS 30µm Cell Strainer (Miltenyi Biotec Ltd 130-041-407) twice. The single cell suspension was centrifuged at 400 x g for 5 mins and resuspended in 0.04% BSA for counting. The cell suspension was placed on ice when possible Cell suspensions containing 17,400 viable cells (aiming for recovery of 10,000 cells) were processed by the Chromium Controller (10x Genomics) and barcoded libraries constructed using the Chromium Next GEM Single Cell 5’ v2 kit (PN-1000263) following manufacturer’s instructions. Samples were sequenced with Illumina HiSeq and reads were mapped and UMI count matrices generated with 10x Genomics CellRanger v.6.0.1 using default parameters and reference transcriptome v. GRCh38-2020. Singlet donor and doublet identity was inferred with cellSNP-lite v. 1.2.0 followed by Vireo v. 0.2.1 (27,28). Single cell data was analysed with Seurat v. 4.0.0 in R v. 4.0.3 (29). Briefly, cells with over 1000 genes, UMI counts under 40,000, under 10% mitochondrial genes, and that had singlet identity predicted by Vireo were retained (doublets ranged from 7-13% of all cells). Cell cycle information for every cell was calculated using Seurat’s *CellCycleScoring()* with marker genes from Tirosh *et al.* (30). Libraries were normalised with *SCTransform()*, and clustering and dimensionality reduction was performed using UMAP within Seurat following the standard pipeline on the merged libraries. Pairwise differentially expressed genes per media contrast were identified with *FindMarkers()* using the Wilcoxon test on log-normalised data, and p-values were adjusted using Bonferroni correction. Cellular identity was inferred by automatic cell type annotation per media sample using singleR (31), which uses a scoring metric for each cell based on correlation of gene expression between the test and the train dataset, using genes that are differentially expressed between the training data labels. The datasets used for annotation were human iPSC-derived microglia matured in mouse xenografts (16) and human foetal macrophage precursors (32). Figures were made with tidyverse (33) and SCpubr (34).

### Identification of perivascular macrophage markers

<u>Publically available data</u> from 1.3 million mouse brain cells was processed as follows (16,32): from a subsample of 20k cells, markers of the microglial/macrophage subpopulation were identified by *FindAllMarkers()* in Seurat; then, a microglial/macrophage identity classifier was built from the top 10 microglial markers with *AddModuleScore()*, which was then applied to score the whole population of 1.3 million cells; cells were considered microglia or macrophages if the score was > 1 (∼5k cells). After normalisation and clustering, three separate clusters of brain macrophage cells (microglia, perivascular macrophages, and macrophage/monocytes) were identified and classified according to the presence of pre-existing markers in the literature in differentially expressed genes calculated via *FindMarkers()*, and converted to human orthologs (**Table S6**). A subset of perivascular macrophage markers (*F13A1*, *LYVE1*, *COLEC12* and *CD163*) were selected for qPCR according to fold expression changes, fraction of microglial cells that express them vs perivascular macrophages, and expression in primary microglia and other reference datasets (data not shown).

### Generation of double fluorescent SH-SY5Y cell line for measuring phagocytic uptake and acidification by microglia

Generation of the SH-SY5Y cell line that reports on its phagocytic uptake and subsequent acidification by microglia (efferocytosis), was as follows. The mCherry-EGFP fragment of pBABE-puro mCherry-EGFP-LC3B (Addgene 22418) was PCR amplified and cloned into a lentivirus backbone (Ef1αF IRES Puro) downstream of EF1α the promotor, using SpeI and BamHI cloning sites (**Figure S1a**). This generated the construct EF1α pmChGIP IRES Puro (pmChGIP, **Figure S1b**) (plasmid available on request). Dual expression of mCherry-eGFP in pmChGIP was confirmed following transfection into HEK293T cells with FuGENE HD (Promega, E2311) (**Figure S1c)**. pmChGIP lentiviral particles were generated by transfecting 4×10^6^ HEK293T cells in 10cm dishes with 4μg EF1α pmChGIP IRES Puro, 2μg psPAX2 (Addgene 12260), 2μg pMD2.G (Addgene 12259). Lentiviruses were harvested at 48hr and 72hr post transfection before centrifugation at 3000rpm to remove any HEK293T cells followed by filtering through a 0.45μm filter low bind PES, processed lentiviruses were stored in single use aliquots at −80°C. Lentiviral titre was calculated using flow cytometry. P10 SH-SY5Y (ATCC, CRL-2266) neuroblastoma cells were transduced at a multiplicity of infection of 2 or 0.3, and double positive mCherry-eGFP cells were single cell sorted at 24hr post transduction into 96 well plates to generate monocultures (**Figure S1d)**. Cells were allowed to clonally expand for several weeks before passaging and banking. Generated clonal lines underwent quality control using flow cytometry to confirm a uniform population of double positive mCherry-eGFP expression (**Figure S1e/f)**. Clone F4 2 was used for assays, due to its high fluorochrome expression. Lines are known as pmChGIP SH-SY5Y and are available upon request.

### Phagocytosis Assays

Microglial precursors were plated at 8.8×10^5^ cells per well in 48 well plates in either the newly optimised medium (ITMG, no coating) or the previous medium (IGBN, Geltrex coating (Haenseler et al)), and matured to microglia for 14d. For imaging, cells were stained for 1 hour at 37°C/5% CO_2_ with 1μM CellTracker Deep Red (Invitrogen, C34565) and 1 drop/mL NucBlue Live ReadyProbes Reagent (Invitrogen, R37605). Cells were washed with PBS, and then incubated for 2hr at 37°C/5% CO_2_ with phagocytic cargo. The three cargo were prepared as follows: 1) Amyloid β aggregates were generated by spiking fluorescent labelled β-Amyloid (Beta-Amyloid (1-42) HiLyte Fluor 488, Anaspec, AS-60479-01) 1:10 with unlabelled β-Amyloid (Beta-Amyloid (1-42) Aggregation Kit, rPeptide, A-1170-1), with shaking incubation for 2 days at 37°C. Aggregates were stored at −80°C and added at a final concentration of 3µM; 2) 3µm carboxylated silica beads (Kisker Biotech, PSI-3.0COOH) were washed in PBS before resuspension in 25mg/mL cyanamide for 15 minutes on a rotator. Beads were washed twice with coupling buffer (0.1M Sodium tetraborate, ddH_2_O, pH 8.0) before labelling for 1 hour with 50µg/mL AF-488 NHS Ester (ThermoFisher, A20000) in coupling buffer. Beads were washed 3 times with quenching buffer (250mM glycine, PBS, pH 7.2) before storage in 0.02% NaN_3_ PBS at 4°C. Beads were added at a 2:1 ratio beads:microglia; 3) pmChGIP SH-SY5Y neuroblastoma cells were maintained in 10% FBS, DMEM/F-12 media at 37°C/5% CO_2_. Cells were harvested with TrypLE Express (Gibco), centrifuged at 400xg for 5 min and resuspended in Hank’s Balanced Salt Solution (HBSS, Lonza). Paraformaldehyde (PFA) (Alfa Aesar) was added to a final concentration of 2% and fixed for 10 min at room temperature. PFA was then diluted in HBSS before centrifugation at 1200xg for 5 min, the cell pellet was washed twice with HBSS. SH-SY5Y cells were added at a 2:1 ratio SH-SY5Y:microglia. Following incubation, iPSC-microglia were washed with PBS to remove non-phagocytosed cargo. iPSC-microglia were live imaged on an EVOS microscope (ThermoFisher), image analysis was undertaken in ImageJ. For flow cytometry, iPSC-microglia were lifted with TrypLE Express (Gibco) and fixed in 2% at room temperature for 10 min. Cells were pelleted at 400xg for 5 min and washed with PBS twice before flow cytometry (Beckman Coulter CytoFLEX Flow Cytometer). Analysis of phagocytic ability was undertaken using FCS Express.

## Results

### IL-34, TGF-β1, M-CSF, and GM-CSF differentially influence morphology and gene expression in iPSC-microglia in monoculture

The previous medium developed by our lab for culturing iPSC-microglia had been optimised for co-culture with neurons, as a result it contained several components that might not be required or optimal for microglial monoculture, including N-2 supplement, β-mercaptoethanol and Geltrex substrate (10). It also did not contain factors that would be expected to be present in co-culture but which might enhance microglial identity if supplemented into monoculture medium - specifically, TGF-β1 (produced by astrocytes and also by microglia (35,36)) and CX3CL1/CD200 (produced predominantly by neurons (37,38)). These factors have been used in some other microglial protocols (12,17). We set out to identify whether removal of neuron-relevant components or addition of these additional factors to the media resulted in differential morphological or transcriptional changes to monocultured iPSC-microglia. In differentiation experimental series 1 (Figure 1b), we matured microglial precursors to microglia for 14 days using 15 different combinations of factors **(Table S4)**, including the original Haenseler *et al.* (10) microglia monoculture medium as the baseline condition (IGBN), the modification of that protocol used by Brownjohn *et al*. (11), (IGF) and also the widely adopted protocol that includes TGF-β1/CX3CL1/CD200 by McQuade *et al.* (ITMCBN) (12).

Visual inspection was used as an initial triage of the 15 conditions (**Figure 2a**). IL-34 as the sole cytokine (I and IBN) was not sufficient for microglial survival, despite its use at 100ng/mL and was therefore not taken forward as sole cytokine. However, co-supplementation with low-dose GM-CSF (IG) or supplementation of IL-34 with M-CSF (MG) supported survival. Removal of either N-2 (IGB) or β-mercaptoethanol (IGN) or both (IG) had no effect on microglial morphology or survival compared with IGBN, consistent with their role in neuronal support rather than for microglia in the iPSC-microglia/neuronal co-culture protocol (10). TGF-β1 induced a clear difference in morphology, microglia were less adherent and more rounded, regardless of further supplementation with other factors (ITG/ITGC/ITGBN/ITGCBN/ITMCBN). CX3CL1/CD200 addition for the final four days of differentiation, as described by (12,16), had no effect on morphology (ITGC/ITGBN/ITGCBN/ITMCBN). FBS (IGF) induced expansion of fibroblasts in the culture, therefore this condition was discarded from future experiments (**Figure 2a**).

**Figure 2:**
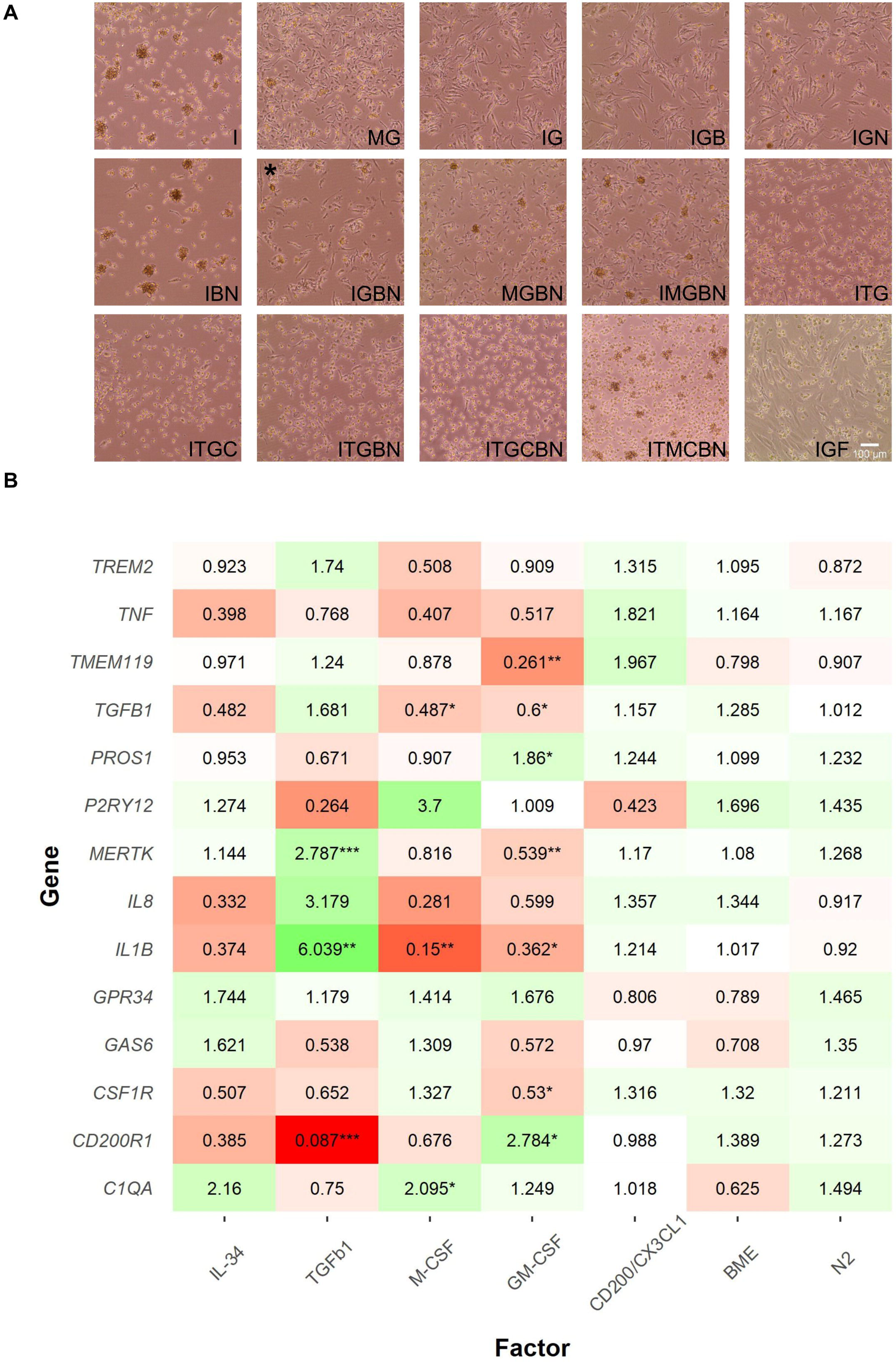
IL-34, TGF-β1, M-CSF, and GM-CSF affect iPSC-microglia morphology and result in transcriptional changes. **A)** Morphology of microglia cultured for 14d in 15 different media combinations on Geltrex. Each letter corresponds to a growth factor or compound; I; IL-34, T; TGF-β1, M; M-CSF, G; GM-CSF, C; CX3CL1/CD200, B; β-mercaptoethanol, N; N-2 supplement, F; Foetal Bovine Serum. Phase contrast images taken at 10x magnification, scale bar 100µm. * media IGBN is media published by Haenseler et al, and is the baseline for comparison. **B)** qPCR results of known microglial marker genes in medium containing different growth factors. Fold changes initially calculated to IGBN before undertaking linear regression, regressing out each individual factor. TGF-β1, M-CSF, GM-CSF all result in transcriptional changes. CD200/CX3CL1, β-mercaptoethanol, and N-2 supplement result in no significant change in microglial gene expression. Fold changes are shown by colour, where green is increased expression and red is decreased and the number within cells. Stars indicate Bonferroni corrected significance, *** p<0.001, ** p<0.01, *p<0.05.

qPCR analysis of microglial markers was consistent with the findings of the morphology observations. Fold changes were calculated to condition IGBN, followed by linear regression (see **Methods**) to examine the effects of inclusion of the different factors (**Figure 2b**). Removal of β-mercaptoethanol or N-2 resulted in no significant differences in gene expression compared to medium where each were included, confirming that β-mercaptoethanol and N-2 have no effect on microglial identity in monoculture. Equally, supplementation with CX3CL1/CD200 for the final three days resulted in no strong transcriptional changes, and were therefore excluded from further experiments. Inclusion or removal of IL-34 resulted in no significant changes to gene expression across all the media conditions versus the baseline condition IGBN, however only two of the fifteen conditions did not include IL-34. M-CSF supplementation resulted in a significant decrease in *IL-1*β (FC, 0.15) and *TGF-*β*1* (FC, 0.49), with *C1QA* (FC, 2.01) significantly increased. TGF-β1 supplementation resulted in a significant decrease in *CD200R1* (FC, 0.09) expression and significant increases in *IL-1*β (FC, 6.04) and *MERTK* (FC, 2.79). Medium in which GM-CSF was included resulted in a significant decrease in expression of *TMEM119* (Fold Change (FC), 0.26), *MERTK* (FC, 0.54), *IL-1*β (FC, 0.36), *CSF1R* (FC, 0.53) and an increase in *PROS1* (FC, 1.86) and *CD200R1* (FC, 2.78).

Combining both the morphology and qPCR data from these initial 15 conditions we concluded that while IL-34, TGF-β1, M-CSF, and GM-CSF result in differential microglial phenotypes, CX3CL1/CD200, β-mercaptoethanol and N-2, have no effect in this experimental setup, and FBS is clearly detrimental.

### Supplementation with TGF-β1 promotes microglial-like identity, whereas M-CSF promotes macrophage identity, these changes are conserved regardless of culturing matrix

Next we sought to identify if supplementation with M-CSF, TGF-β1, or a combination of both would support microglial differentiation compared to our original baseline condition, IGBN medium. In parallel, we tested different tissue culture matrices that have been used in published protocols (10,12,17).

We trialled five media conditions in differentiation two with combinations of factors identified by our initial triage: the baseline condition IGBN (Haenseler et al 2017) (IL-34, low-dose GM-CSF, β-mercaptoethanol, N-2); IMBN (IL-34, M-CSF, β-mercaptoethanol, N-2), where GM-CSF is replaced with M-CSF; ITGBN (IL-34, TGF-β1, GM-CSF, β-mercaptoethanol, N-2) where the baseline IGBN condition is supplemented with TGF-β1; ITM RPMI (IL-34, TGF-β1, M-CSF) a medium previously described by (16) and ITM ADMEM (IL-34, TGF-β1, M-CSF, Advanced DMEM basal media). ITM ADMEM ensured the same basal media was used for all factor cocktails. We plated cells at the same density on three different matrices: standard TC treated plastic; Geltrex coated or 10μg/ml fibronectin coated.

iPSC-microglia cultured on tissue-culture plastic were observed to have greater attachment and confluency than on Geltrex, which was greater than fibronectin **(Figure 3a)**. There was no notable difference in morphology between IGBN, IMBN, and ITGBN on any of the different matrices. However, ITM resulted in poor attachment, especially in RPMI base, and this was only partly rescued by ADMEM base in combination with TC plastic. The minimal composition of RPMI media versus ADMEM may play a role here.

**Figure 3:**
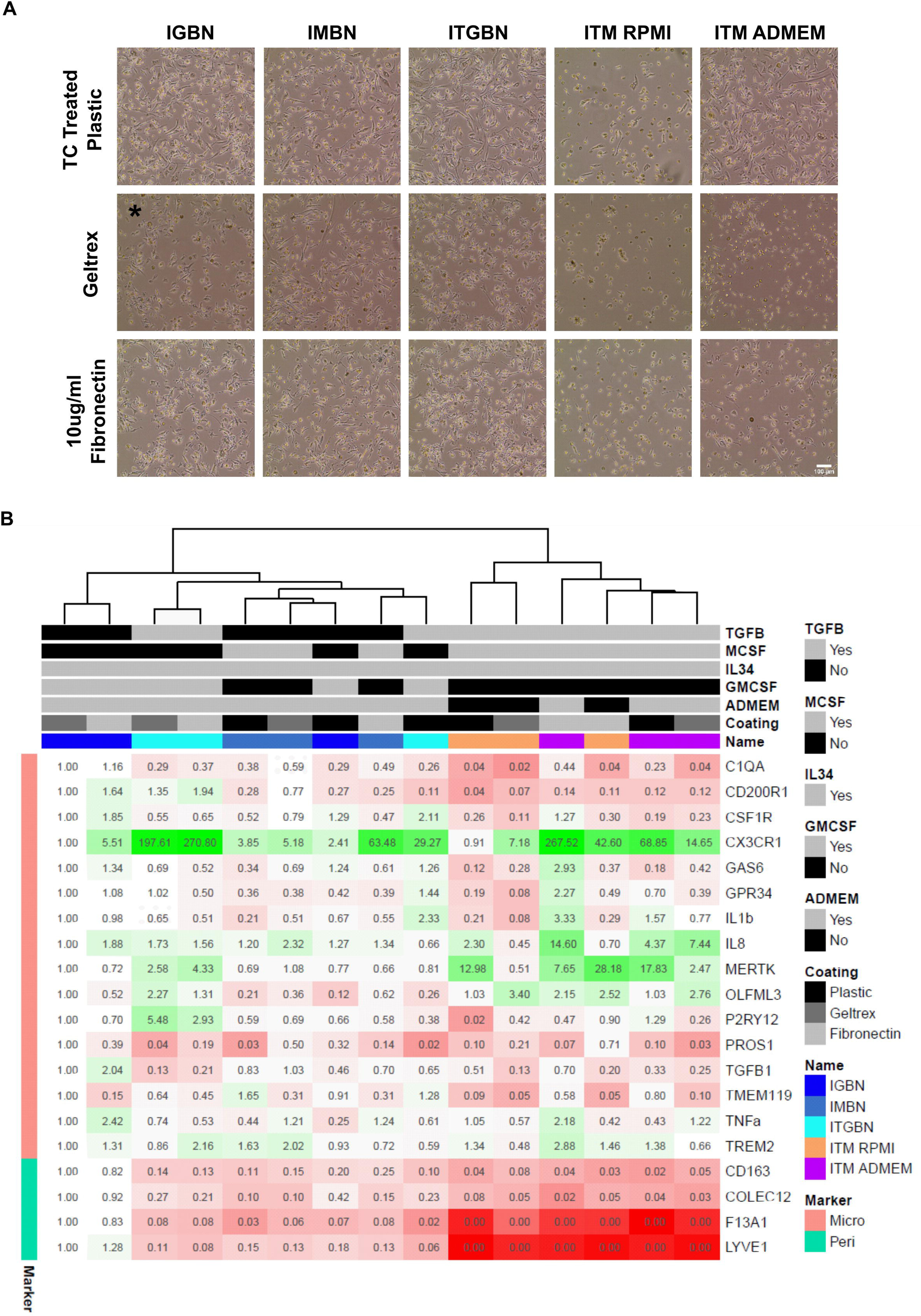
TGF-β1 promotes microglial identity, and M-CSF promotes macrophage identity, tissue culture coating influences morphology but medium factors are the main driver of identity. **A)** Morphology of iPSC-microglia cultured for 14d in five different media combinations on either tissue culture treated plastic, Geltrex, or 10µg/ml fibronectin. Each letter corresponds to a growth factor or compound; I; IL-34, T; TGF-β1, M; M-CSF, G; GM-CSF, B; β-mercaptoethanol, N; N-2 supplement.. Phase contrast images taken at 10x magnification, scale bar 100µm. * media IGBN is media published by Haenseler et al, and is the baseline for comparison. Geltrex resulted in the fewest cells per media condition, followed by fibronectin, with TC treated plastic resulted in the highest cell number. ITM with RPMI base medium results in a large reduction of cell numbers compared to the medium with ADMEM/F12 base. **B)** qPCR of the 15 samples in A) for microglial and perivascular macrophage markers, fold changes represented relative to IGBN Geltrex (the Haenseler et al protocol). Fold changes are presented as both colour changes (green indicates increased expression, red decrease expression) and the number within each cell. Samples cluster by medium, with medium containing TGF-β1 showing lower expression of perivascular macrophage genes (*F13A1/LYVE1/COLEC12)* and increased microglial genes (*CX3CR1/MERTK/OLFML3)*.

qPCR analysis identified that medium composition was the main driver of clustering, resulting in much larger transcriptional changes than tissue culture coating (**Figure 3b**). For this qPCR panel we included perivascular macrophage markers (*F13A1/LYVE1/COLEC12/CD163*), as identified by differential expression of brain macrophage subpopulations (see **Methods**), and which have been described previously in the literature (39,40). These were highest within the baseline IGBN media and were downregulated when the media was supplemented with TGF-β1. Conversely, TGF-β1 supplementation (ITGBN) resulted in increased expression of microglial identity genes *CX3CR1* (29/197/270 fold increase on plastic, Geltrex, and fibronectin respectively), *MERTK* (0.8/2.6/4.3), *OLFM3* (0.3/2.3/1.3) and *P2RY12* (0.4/5.5/2.9). Note also that these microglial identity genes all are increased when cells are cultured on a matrix. Replacing GM-CSF with M-CSF (IGBN vs IMBN) reduced expression of perivascular macrophage markers. Microglial markers (*CX3CR1/MERTK/OLFML3*) were also increased in IMBN, however the fold changes were lower than for ITGBN, indicating that TGF-β1 is the key driver in microglial identity.

Interestingly, co-supplementation of TGF-β1 with M-CSF resulted in the strongest microglial identity signal (ITM), with *MERTK, IL8, and OLFML3* showing the largest fold change (*CX3CR1* was also upregulated but the changes were not as great as ITGBN). Basal media (ADMEM/F12 vs RPMI ATCC) had little effect on the gene expression profile of the microglia.

These gene expression profiles correlated well between independent differentiations at two independent sites, using iPSCs with different genetic backgrounds (**Figure S2**), indicating that the media conditions tested result in robust changes in gene expression (IMBN, R^2^=0.236, p-value=0.03. ITGBN, R^2^=0.348, p-value=0.006). Given the close correlation in gene expression between ITGBN and ITM we next assessed whether inclusion of low-dose GM-CSF would improve cell retention while maintaining microglial identity.

### Low-dose GM-CSF promotes cellular retention while maintaining microglial identity

Our final differentiation series (differentiation three, **Figure 1b**) examined the effect of low-dose GM-CSF (10ng/ml) on the retention of iPSC-microglia in monoculture We assessed six conditions: the baseline IGBN medium (10); modified ITM ADMEM medium (from (17)); and ITM supplemented with GM-CSF (ITMG), each on tissue culture treated plastic versus fibronectin-coating, then undertook qPCR for microglial and perivascular marker gene expression.

Tissue culture treated plastic versus fibronectin coating had no effect on morphology across all three media (**Figure 4a**). ITM ADMEM medium resulted in decreased adherence and a more rounded morphology compared to IGBN/ITMG, in agreement with the previous experimental series. Supplementation with low-dose GM-CSF (ITMG, versus ITM) resulted in stronger retention on the substrate and resultant increased yield of cells at the end of the 14 days of culture, this is supported by previous work in neutrophils and macrophages (41,42).

**Figure 4:**
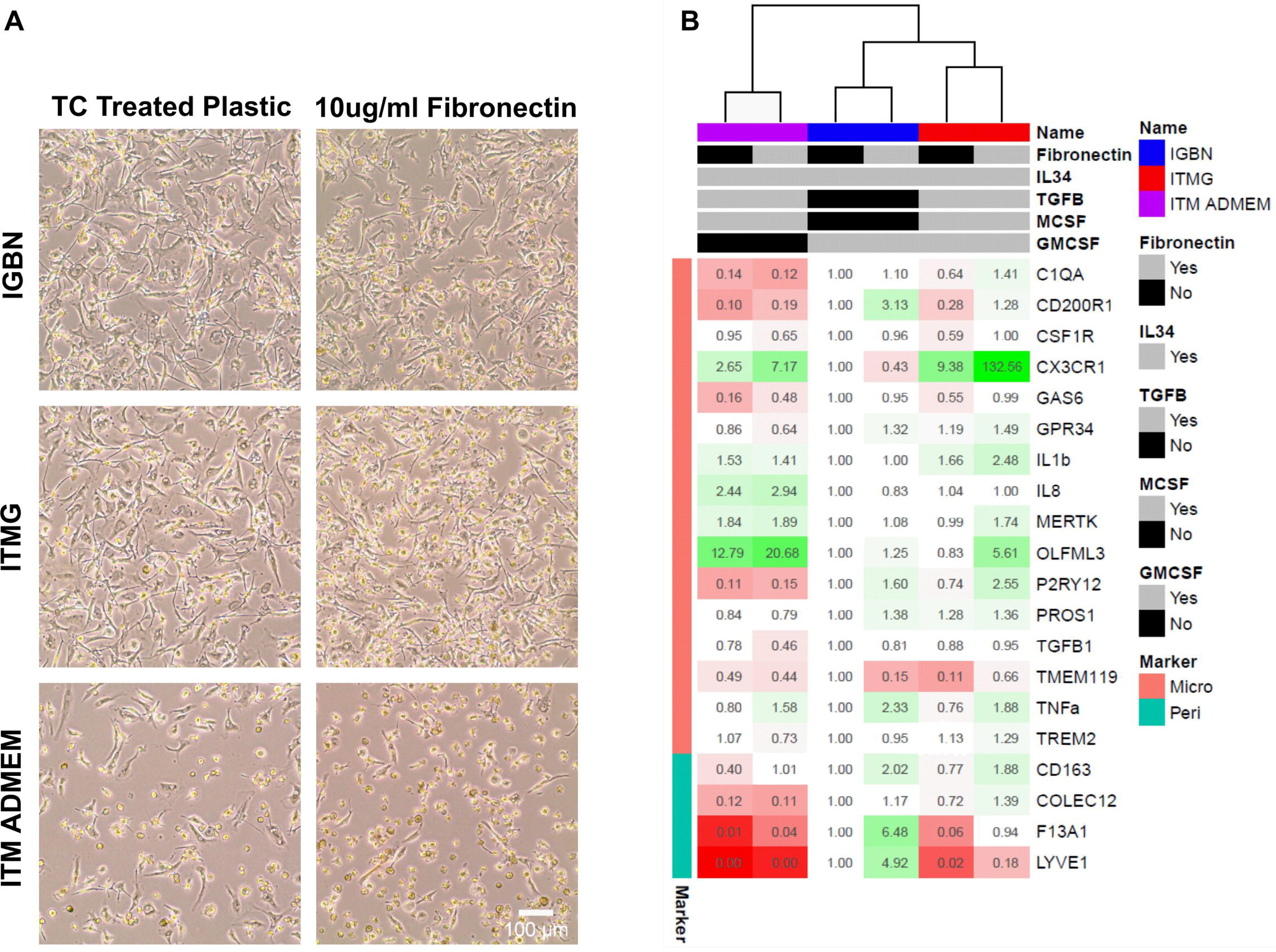
Low level supplementation with GM-CSF promotes cellular survival while maintaining microglial identity. **A)** Morphology of microglia cultured for 14d in three different media combinations on either tissue culture treated plastic or 10µg/ml fibronectin. Each letter corresponds to a growth factor or compound; I; IL-34, T; TGF-β1, M; M-CSF, G; GM-CSF, B; β-mercaptoethanol, N; N-2 supplement. Phase contrast images taken at 10x magnification, scale bar 100µm. Media IGBN (Haenseler et al 2017) is the baseline for comparison. ITM ADMEM results in low survival and adherence on both TC treated plastic and 10µg/ml fibronectin, however when supplemented with GM-CSF (ITMG) the survival and adherence is improved. **B)** qPCR of the six samples from A) for microglial and perivascular macrophage markers, fold changes represented as changes to IGBN TC treated plastic. Fold changes are presented as both colour changes (green indicates increased expression, red decrease expression) and the number within each cell. Samples cluster by medium followed by coating. ITMG promotes microglial identity with increased expression of *CX3CR1* and *OLFM3* and a reduction in perivascular macrophage markers (*LYVE1/F13A1)*. Coating with fibronectin results in an increased expression of microglial identity genes in ITM and ITMG.

qPCR indicated strong clustering by media rather than by coating (**Figure 4b**), in accordance with the previous experimental series. As previously, inclusion of TGF-β1 decreased perivascular marker expression (*COLEC12/LYVE1/F13A1*), and increased standard microglial marker gene expression (*CX3CR1*, *OLFML3*, *MERTK)*. Microglia cultured in ITM ADMEM had decreased expression of *CD200R1, P2RY12* and *C1QA,* but increased *OLFML3* versus IGBN/ITMG. The strongest microglial identity signal was in iPSC-microglia cultured in ITMG on fibronectin. Again, these results were highly correlated between two independent labs, independent differentiations, and different genetic backgrounds (ITM, R^2^=0.639, p-value=2.36×10^−5^, ITMG, R^2^ =0.422, p-value=0.00195) (**Figure S3, S4)**.

While the panel of qPCR markers helped to confirm microglial identity, this was limited to a few select marker genes, so we next explored whether the different media result in different subtypes of microglia (43,44) by undertaking single cell RNAseq on a subset of conditions.

### Single cell RNA-seq confirms that TGF-**β**1 promotes microglial identity, while medium containing M-CSF only promotes macrophage identity

To get a more detailed picture of the transcriptional landscape of iPSC-microglia generated by the different media, we undertook scRNA-seq of cells cultured in the baseline medium IGBN, and IM, IMBN, ITGBN, ITM ADMEM and ITMG, all on TC plastic. After quality control, filtering of doublets and low-quality cells, normalization and clustering (see **Methods, Figure 5a, S5a**), we could infer higher transcriptomic similarity between ITM ADMEM and ITMG media. The examination of cell marker distribution indicated some heterogeneity in microglial marker expression among the different media, with reduced density of perivascular marker gene expression in ITM ADMEM and ITMG (**Figure S5b**). Differential expression analysis of each medium against the IGBN baseline resulted in a mean of 1196 significant differentially expressed (DE) genes, with the ITM ADMEM and ITMG comparisons resulting in the largest amount of DE genes (**Figure S5c, Table S7**). These DE genes included many of the previously highlighted microglia and perivascular macrophage markers (**Figure 5b**). Analysis of microglial and perivascular markers for these media showed again a significant increase in the expression of the microglial genes in media including TGF-β1, and the highest perivascular macrophage gene expression in IGBN, IM and IMBN media.

**Figure 5:**
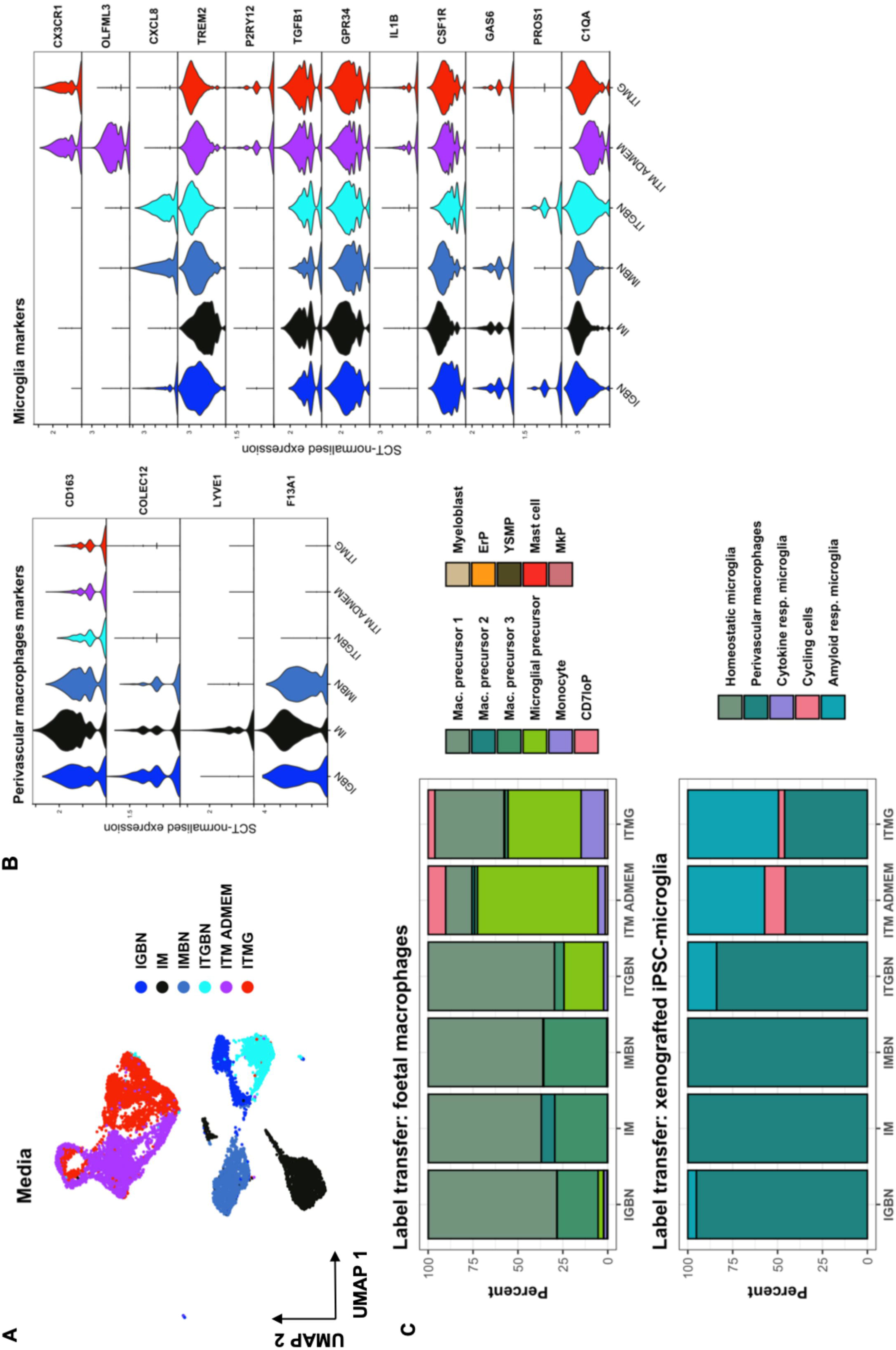
Single cell RNA-seq confirms that TGF-β1 promotes microglial identity. **A)** Uniform manifold approximation and projection (UMAP) plot of quality-controlled iPSC-derived microglia cells, coloured by differentiation media. **B)** Violin plots of SCT-normalised expression (y-axis) per media (x-axis) per perivascular macrophage or microglia marker gene. **C)** Label transfer with singleR: cumulative percentage of cells (y-axis) of each label (colour) per media (x-axis). Mac. precursor 1-3; macrophage precursors subtypes, CD7loP; CD7lo progenitor, ErP; erythroid progenitor, YSMP; yolk sac-derived myeloid-biased progenitor, MkP; megakaryocyte progenitor, Cytokine / Amyloid resp. microglia; Cytokine / Amyloid response microglia.

We then expanded our investigation of the transcriptional identity of these cells to the whole genome by performing unbiased cell type annotation (“label transfer”) with singleR using two reference datasets. The first one comprises of foetal macrophage precursors and other hematopoietic cells isolated from human embryos (32), including developing microglia, and was chosen due to the perceived similarities between iPSC-derived cells and immature microglia. The second dataset is comprised of iPSC-derived microglia whose last step of maturation has been performed as xenografts in mouse brain (16), and represents the *in vitro* – generated microglia that would be most similar transcriptomically to human primary microglia. Label transfer results confirm the increased microglial identity when using media including TGF-β1 (**Figure 5c**, **S5d**), particularly ITM ADMEM and ITMG, as denoted by the larger proportion of cells annotated as “microglial precursors” in the foetal macrophage dataset, and the reduced number of perivascular macrophages as classified by the xenografted iPSC-microglia dataset.

While cells matured in ITM ADMEM and ITMG are both highly classified as microglia, the reduced reproducibility and poorer retention of cells cultured in ITM across different differentiations led us to conclude that ITMG is preferable, as it not only has a similar transcriptional profile to ITM ADMEM but also has improved cell retention and reproducibility across differentiations. Therefore ITMG was taken forward for final functional characterisation versus our original IGBN medium.

### Microglia differentiated in optimised ITMG medium exhibit phagocytic competence

Finally we examined the ability of microglia matured in the newly optimised ITMG medium to take up phagocytic cargo, compared to the baseline medium, IGBN. We utilised three different cargos: fluorescent Amyloid-β aggregates; 488-labelled silica beads; and a novel double-fluorescent SH-SY5Y cell line to measure dead neuron uptake and acidification in the phagolysosome. The double fluorescent SH-SY5Y cell line expresses a GFP-mCherry fusion protein which becomes single mCherry fluorescent when phagocytosed into the low pH environment of the phagolysosome due to pH sensitive conformational changes resulting in loss of GFP fluorescence. As expected, different cargos were taken up by the cells at different rates (Amyloid-β >silica beads >dead SH-SY5Y) (**Figure 6**). Importantly, regardless of the cargo, we observed no significant difference between microglia cultured in IGBN versus ITMG media, demonstrating that the improved medium does not alter this important microglial function.

**Figure 6:**
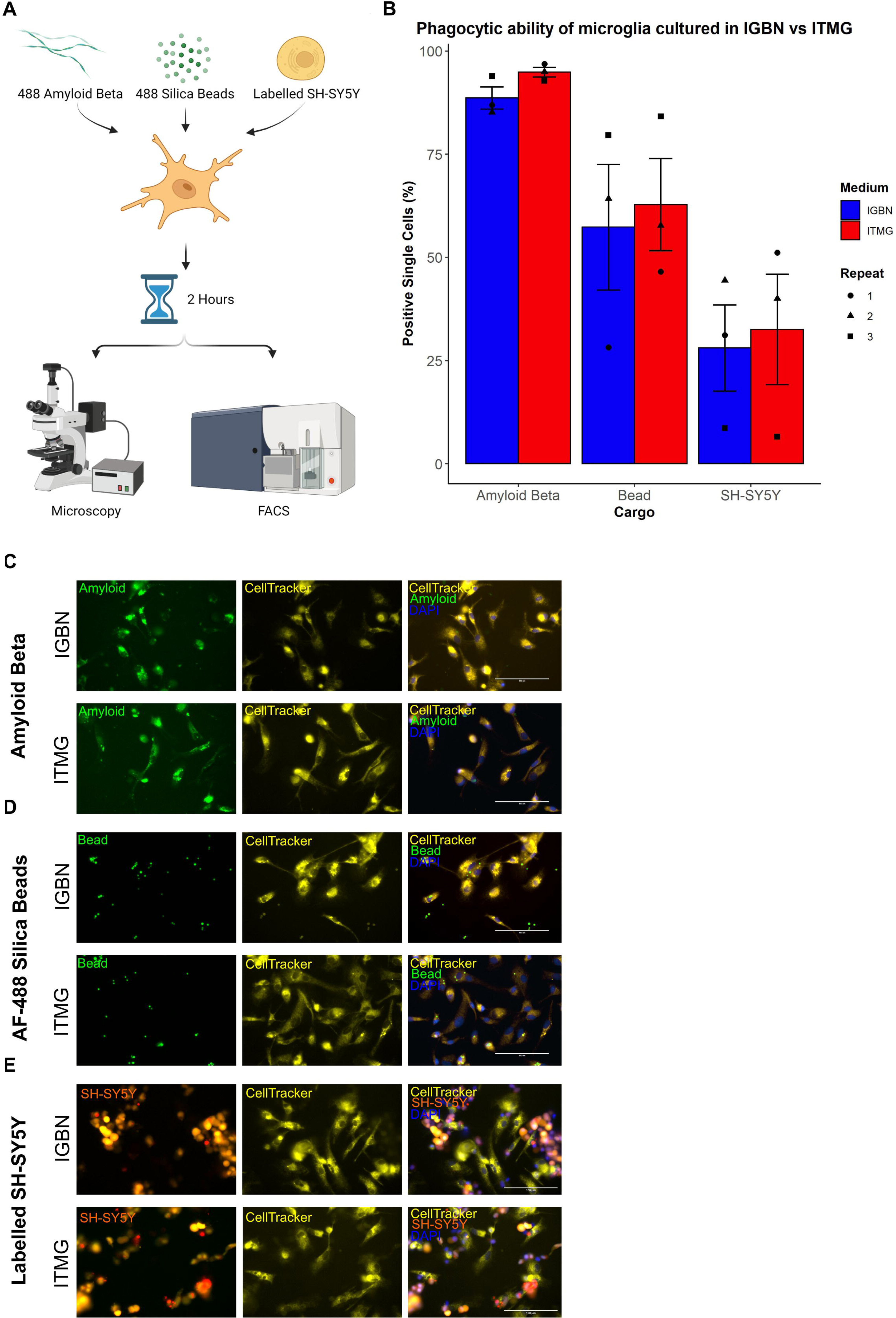
Functional analysis of phagocytosis results in no difference between the new ITMG medium and our previously published IGBN medium. **A)** To examine phagocytosis in differentiated iPSC-microglia in our new medium, we incubated cells with three different cargos; Amyloid-β aggregates, 488 labelled silica beads, and fluorescent labelled dead SH-SH5Y neuroblastoma. Phagocytosis occurred for 2 hours before fixing the microglia and quantifying fluorescence using FACS, and representative images taken by microscopy. **B)** There is a difference in the phagocytic ability of the different cargos, however there is no difference between the two medium. Data presented as mean ±SEM, N=3 independent replicates, with individual data points shown. Each independent replicate contained three technical repeats. **C)** Representative images of Amyloid-β phagocytosis at two hours. **D)** Representative images of AF-488 labelled bead phagocytosis at two hours. **E)** Representative images of labelled dead SH-SY5Y neuroblastoma phagocytosis is two hours, red indicates a SH-SY5Y which has been phagocytosed and is in low pH environment. All images are shown at 40x magnification, microglia are marked with CellTracker, and nuclei are stained with DAPI.

## Discussion

Multiple papers over the last five years have described different methods for differentiating iPSC to microglia, resulting in variation between labs and potential reproducibility issues when translated to different institutes or genetic backgrounds. For iPSC-microglia in monoculture the field has not yet fully consolidated which factors are crucial for the development of iPSC-microglia and some media contain redundant “legacy” factors, their role having never been assessed systematically. Moreover, some protocols use poorly defined components, including conditioned media (45). We set out to systematically identify the key defined factors required for microglial identity in monoculture, utilising morphology, qPCR for both microglial and non-microglial brain macrophages, and scRNA-seq, across two laboratories and multiple genetic backgrounds, to provide a highly reproducible differentiation and single-cell level characterisation of the resulting microglial population.

We started by identifying the defined components used in various iPSC-microglial protocols and tested in parallel their ability to produce iPSC-microglia in monoculture, compared to our original medium (IGBN). Initial morphological assessment allowed us to exclude medium containing FBS (fibroblast outgrowth) and IL-34-only supplementation (increased cell clumping and death). Media where IL-34 was supplemented with M-CSF or low-dose GM-CSF showed increased survival. M-CSF and IL-34 engage the myeloid receptor CSF1R, and GM-CSF acts on the receptor CSF2R, both of which activate survival signalling pathways (46,47). IL-34 is the main ligand for CSF1R in the brain, whereas M-CSF is the main ligand for CSF1R in the periphery, both mediating a similar intracellular response (46). Supplementation with M-CSF is known to promote a macrophage phenotype and has been used in multiple iPSC-macrophage protocols, and the scRNA-seq analysis here shows that M-CSF-alone induces a more macrophage-like phenotype (21,23,48). This supports a unique role of IL-34 in microglial development in the brain which cannot be replicated by peripheral M-CSF alone (49,50).

qPCR and morphological analysis confirmed that β-mercaptoethanol and N-2 supplement have no effect iPSC-microglial differentiation or transcription so can be included if microglia are to be used in co-culture (10). Most surprisingly was the lack of effect of CX3CL1/CD200 on the phenotype of iPSC-microglia when others have observed changes (16,51). Both CX3CL1/CD200 are immunoregulators expressed by neurons, and ligands for CX3CR1/CD200R on microglia. Nonetheless, the addition of TGF-β1 (another potent immunoregulator) could make the role of CX3CL1/CD200 redundant (37,38,52,53). These initial results reduced our factor list from nine to four (IL-34, TGF-β1, M-CSF, GM-CSF) for further exploration.

We identified consistent upregulation of microglial marker genes *CX3CR1, MERTK and OLFML3* in medium containing TGF-β1. This is unsurprising, as TGF-β1 has been shown to be critical to microglial maturation in mice (54). Furthermore, adult primary microglia cultured in M-CSF and TGF-β1 have been shown to have increased microglial identity versus M-CSF or GM-CSF alone (54), which is replicated here in iPSC-microglia. The scRNA-seq data further supports the role of TGF-β1 in microglial maturation versus M-CSF in macrophage maturation.

The low cell retention and poor reproducibility of microglia matured in ITM led us to explore if GM-CSF would promote cellular retention while maintaining microglial identity. GM-CSF is a cytokine expressed by astrocytes and during neurodevelopment (55,56). GM-CSF has been identified to improve antigen presentation in murine primary microglia (57) and has been known to modify microglial morphology (10,58). This is supported in our findings, where ITMG microglia are more adherent and ramified versus ITM microglia, regardless of culturing matrix, and the phenotype more consistent across differentiations than ITM microglia. Examining the transcriptome through qPCR identified several changes between IGBN and ITMG. Notably, the reduction in the perivascular macrophage markers (*F13A1/LYVE1*) and increase in microglial markers (*CX3CR1/OLFML3/P2RY12)*. The microglial identity is further improved when cells are cultured on fibronectin, while media is the main driver of identity the use of fibronectin is left to the discretion of the user.

## Conclusion

Coupling the results of morphological assessment, qPCR, scRNAseq, and phagocytosis assays, we have demonstrated that the optimised ITMG medium described here improves on our previously published IGBN medium for culturing iPSC-microglia. Differentiation of iPSC to mesoderm and hemogenic endothelium via embryoid bodies, then bulk myeloid differentiation to primitive macrophage/microglia precursors, and final maturation of monoculture iPSC-microglia in ITMG medium, provides a highly scaleable pipeline for routine experiments investigating neuroinflammation.

## Acknowledgements

This work was supported by a research grant from Open Targets (Project OTAR2065).

## Author contributions

S.J.W, Y.C, J.S performed all laboratory experiments. S.J.W, Y.C, J.S, M.P.A analyzed the data. M.P.A provided statistical and computational insight. W.S.J, S.J.W, S.A.C, conceived and planned experiments. S.J.W, M.P.A, Y.C, J.S, W.S.J, S.A.C, A.R.B prepared the manuscript. All authors discussed results and contributed to the final article.

## Data Availability

All single cell RNAseq data is available upon request to Open Targets https://www.opentargets.org/

## Figures and Legends

**Supplementary figure 1: Development of the pmChGIP SH-SY5Y cell line for measuring autophagy. A)** The pmChGIP construct was generated by PCR amplification of mCherry-eGFP fusion from the pBABE-puro-mCherry-eGFP-LC3B vector with SpeI and BamHI restriction sites. These were cloned into a lentiviral backbone EF1α IRES Puro to generate the pmChGIP vector. **B)** Plasmid map of pmChGIP, **C)** HEK293T were transfected with pmChGIP (dual mCherry eGFP vector, pmCherry (single mCherry vector), or pGIP (single eGFP vector) to confirm dual expression. Images taken at 48hr post transfection using EVOS Floid at 20x magnification. Scale bar 100µm. Cont are untransfected HEK293T cells. **D)** SH-SY5Y at 48hr following transduction with pmChGIP lentivirus confirms successful integration and signal. Images taken using EVOS Floid at 20x magnification. Scale bar 100µm. Cont are untransduced SH-SY5Y. **E)** Clonal pmChGIP SH-SY5Y cell lines were generated by transduction of pmChGIP at 1:32 or 1:2 dilution of lentivirus (corresponding to MOI of 0.3 and 2 respectively) into p10 SH-SY5Y. After 24hr single cell sorting was undertaken for double positive cells. Sorted cells were expanded over several weeks before banking. Fluorescence of the four 1:32 pmChGIP SH-SY5Y lines and two 1:2 pmChGIP SH-SY5Y lines were quantified using FACS. Cells were either live or fixed, to mimic the phagocytosis assay conditions. Fixing reduced the fluorescent intensity of all lines as expected. Line pmChGIP 1:2 F4 showed the highest geometric mean intensities and was used for phagocytosis assays. **F)** Example FACS plots showing clear separation of control SH-SY5Y and the pmChGIP 1:2 F4 SH-SY5Y cell line. 488 signal on the X-axis and 561 signal on the Y-axis.

**Supplementary figure 2: Overlapping qPCR profiles of IMBN and ITGBN from two different institutes with different genetic backgrounds confirms reproducibility.** ddCT values for iPSC-microglia cultured in IMBN **A)** and ITGBN **B)** at both Oxford and Cambridge using different genetic backgrounds show a strong correlation in their expression profiles. ddCT values calculated against IGBN. (IMBN, R^2^=0.236, p-value=0.03. ITGBN, R^2^=0.348, p-value=0.006).

**Supplementary figure 3: Overlapping qPCR profiles of ITMG and ITM from two different institutes with different genetic backgrounds confirms reproducibility.** ddCT values for iPSC-microglia cultured in ITMG **A)** and ITM ADMEM **B)** at both Oxford and Cambridge using different genetic backgrounds show a strong correlation in their expression profiles. ddCT values calculated against microglial precursors. (ITM, R^2^=0.639, p-value=2.36×10^−5^, ITMG, R^2^ =0.422, p-value=0.00195)

**Supplementary figure 4: Representative images of iPSC-microglia differentiated for 14d on three different genetic backgrounds (SFC841-03-01, SFC856-03-04, KOLF2.1S).** Images taken at d14 at 10x magnification. Scale bar 100µm.

**Supplementary figure 5: Single cell RNA-seq analysis of six media. A)** UMAP visualizations of cells coloured by clusters or inferred cell cycle phase (top), and principal component analysis (PCA) visualizations of cells coloured and split by media (bottom). **B)** UMAP visualizations of cells coloured by gene expression density of perivascular macrophage and microglial marker genes. **D)** Intersecting set plot of differentially expressed genes between ITGBN, IMBN, IM, ITMG and ITM ADMEM media against the IGBN baseline. Total differentially expressed genes per contrast in “Set Size”. **E)** UMAP visualizations of cells coloured by label transfer assignments per training data.

