## Supplementary figures and images for "Single-cell transcriptomics defines an improved, validated monoculture protocol for differentiation of human iPSCs to microglia"

### Supplementary Figure 1

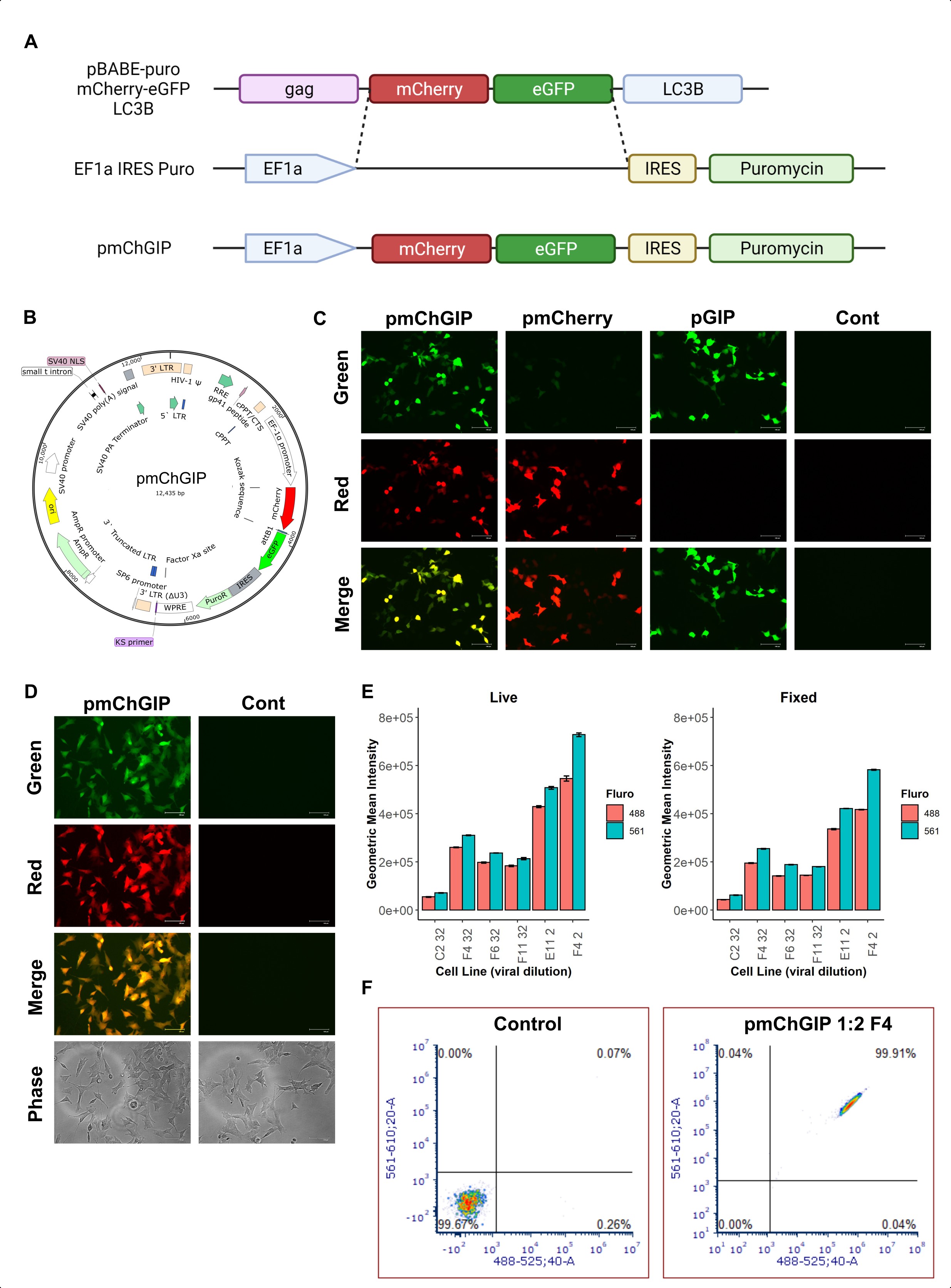

### Supplementary Figure 2

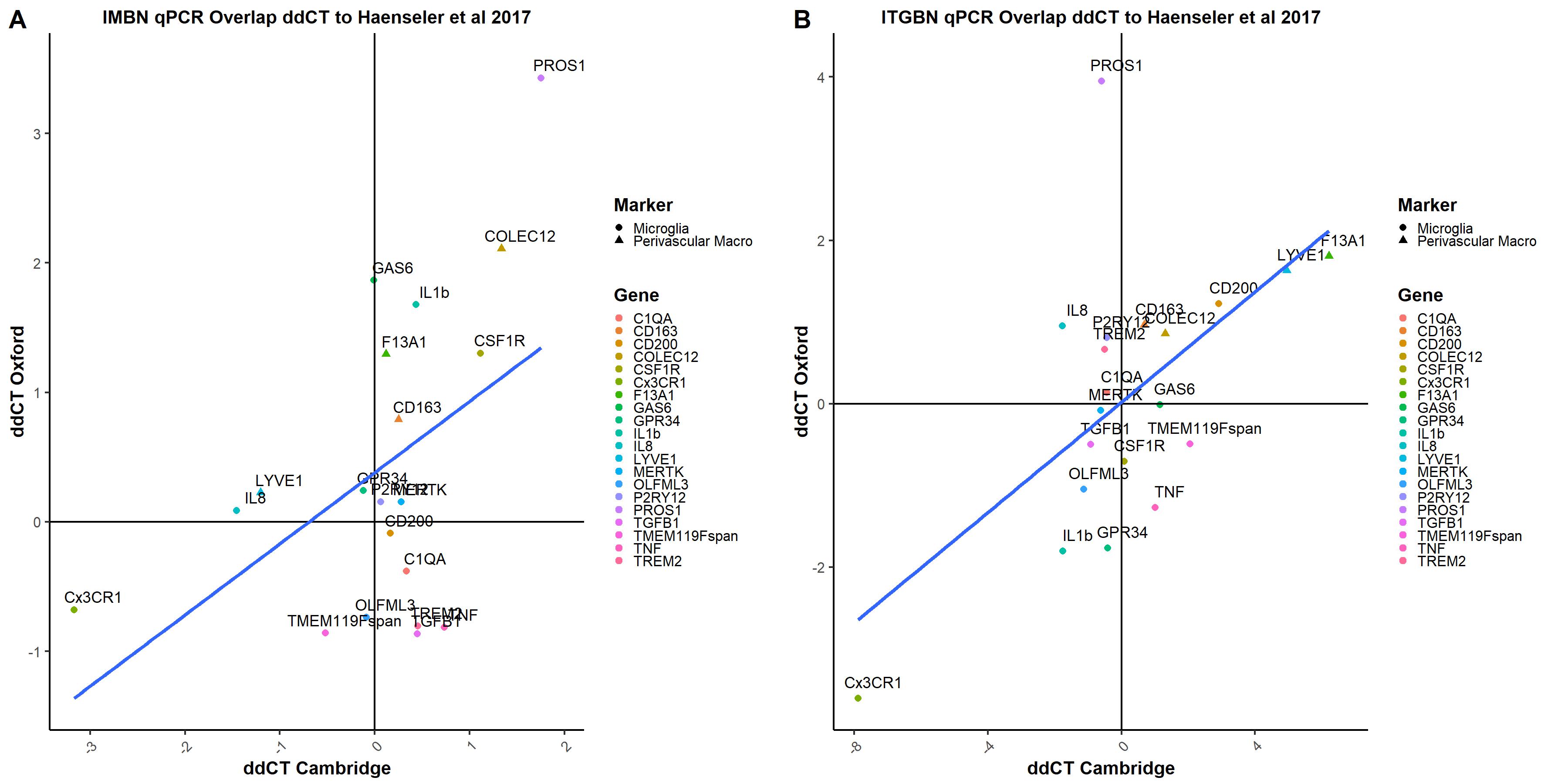

### Supplementary Figure 3

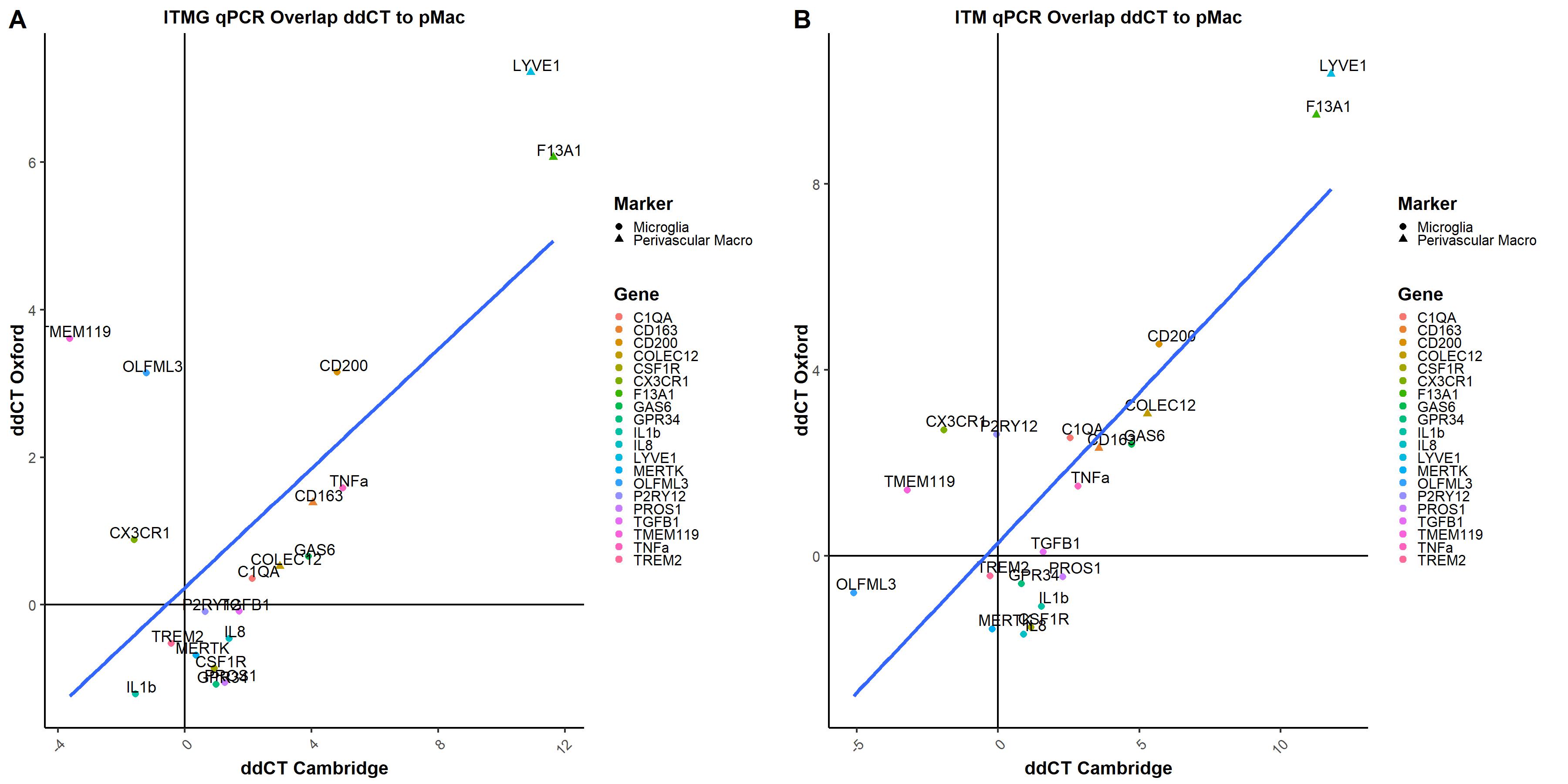

### Supplementary Figure 4

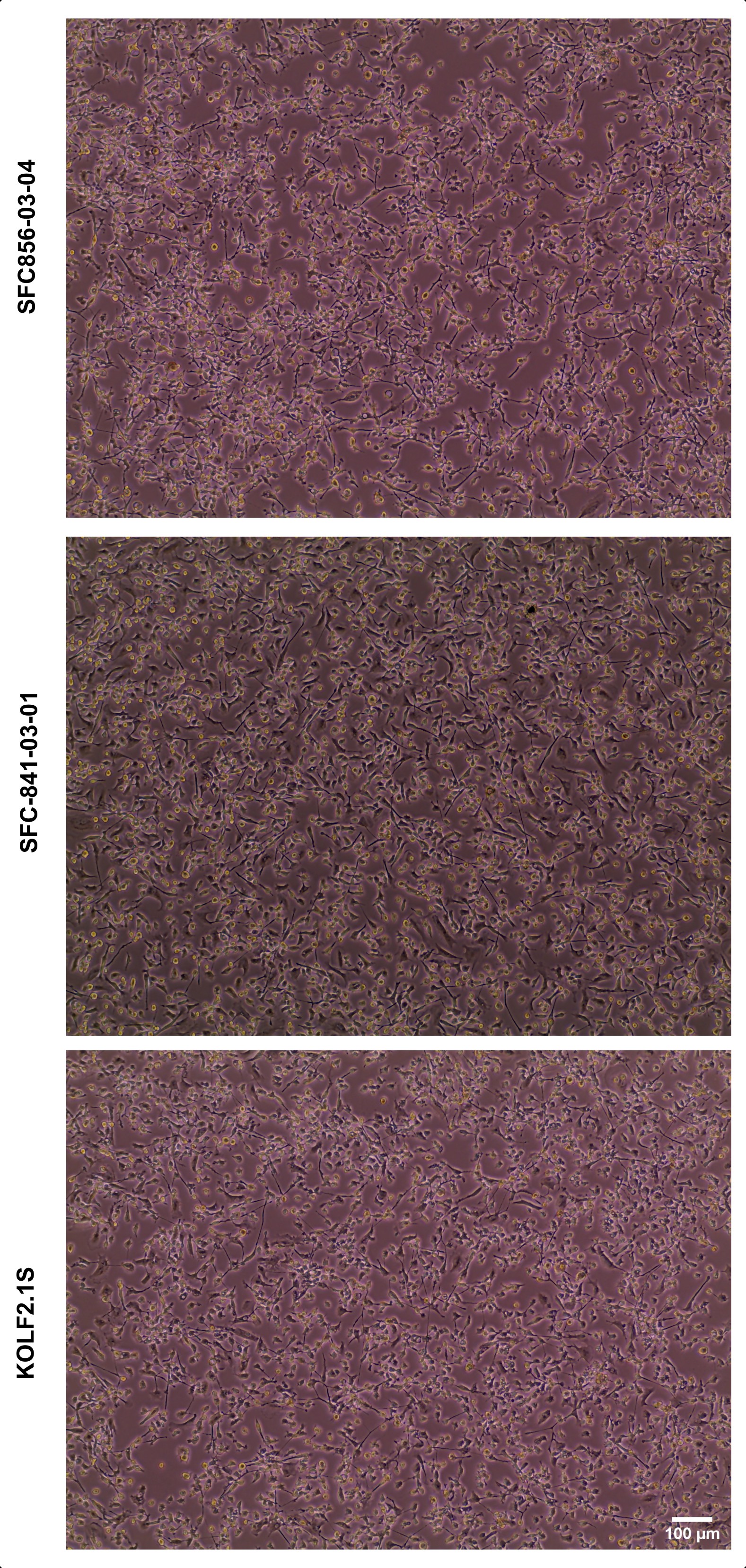

### Supplementary Figure 5

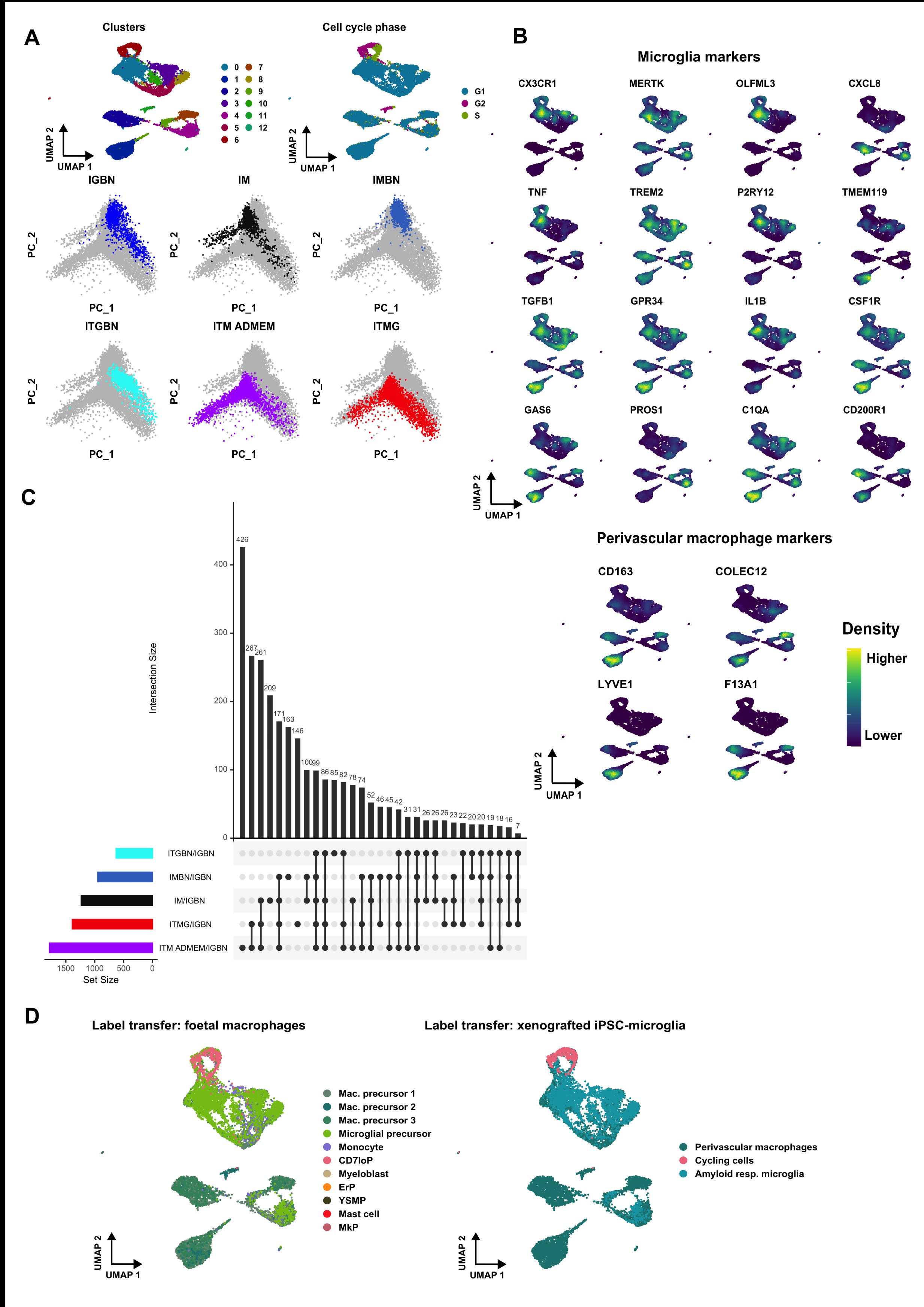
